# Genomic drivers of large B-cell lymphoma resistance to CD19 CAR-T therapy

**DOI:** 10.1101/2021.08.25.457649

**Authors:** Michael D. Jain, Bachisio Ziccheddu, Caroline A. Coughlin, Rawan Faramand, Anthony J. Griswold, Kayla M Reid, Ola Landgren, Frederick L. Locke, Francesco Maura, Marco L. Davila, Jonathan H. Schatz

## Abstract

Chimeric antigen receptor-reprogrammed autologous T cells directed to CD19 are breakthrough immunotherapies for heavily pretreated patients with aggressive B-cell lymphomas but still fail to cure most patients. Host inflammatory and tumor microenvironmental factors associate with CAR-19 resistance, but the tumor-intrinsic factors underlying these phenomena remain undefined. To characterize genomic drivers of resistance, we interrogated whole genome sequencing of 30 tumor samples from 28 uniformly CAR-19-treated large-cell lymphoma patients. We reveal that patterns of genomic complexity (i.e., chromothripsis and APOBEC mutational activity), and distinct genomic alterations (deletions of *RB1* or *RHOA*) associate with more exhausted immune microenvironments and poor outcome after CAR-19 therapy. Strikingly, pretreatment reduced expression or sub-clonal mutation of CD19 did not affect responses, suggesting CAR-19 therapy successes are due not only to direct antigen-dependent cytotoxicity but require surmounting immune exhaustion in tumor microenvironments to permit broader host responses that eliminate tumors.

## INTRODUCTION

Chimeric antigen receptor-modified T cells targeting CD19 (CAR-19) are among new immunotherapy options for patients with diffuse large B-cell lymphoma (DLBCL)^1–3^. Unfortunately, treatment failures and relapses are common^4–9^, and underlying mechanisms remain unclear. Disease aggressiveness and serum inflammatory markers associate with poor outcome^5,10,11^, as does T-cell exhaustion in either the tumor microenvironment (TME)^12^ or the CAR-19 product^13^. Efforts to improve efficacy such as dual-targeting strategies^14–16^ remain uninformed by an understanding of the lymphoma cell-intrinsic factors that drive CAR-19 failures. In particular, there is a lack of knowledge on tumor cell genomic drivers involved in relapse.

We therefore were motivated to dissect the role of genomic drivers and their association with the TME changes that thwart CAR-19 efficacy. We performed the first ever whole-genome sequencing (WGS) analysis of large B-cell lymphoma tumors from patients uniformly treated with the CAR-19 product axicabtagene ciloleucel (axi-cel). We find resistance associated with specific genomic findings including chromothripsis events, apolipoprotein B mRNA-editing enzyme, catalytic polypeptide (APOBEC) mutational activity, point mutations in distinct driver, and deletions of *RB1* or *RHOA. CD19* genomic loss and/or low expression by flow cytometry were mostly confined to patients with complete responses and excellent outcome, and all samples collected at relapse expressed CD19. These data suggest CAR-19 clinical activity is driven not only by the interaction between the engineered immune effector and CD19 but also by promoting a broader immune attack that is more likely to be thwarted by genome complexity and tumor aggressiveness than by loss of the CAR-targeted antigen.

## RESULTS

### Patient cohort

LBCL tumor biopsies (with paired germline samples) of 31 patients treated with axi-cel were analyzed by WGS (median coverage 44.3X, range 30.39-76.08, **Supplementary Table 1**). Of the initial 31 cases, three failed sequencing due to low cancer cell fraction (CCF) and normal match contamination. Most tumors were sampled immediately prior to CAR-19 therapy, with two cases containing relapse-only biopsies and two with both pre-CAR-19 and relapse samples. Demographics, disease characteristics and response to axi-cel treatment for the 28 patients with reportable data are summarized in **Table 1**. All had large B-cell lymphoma – 24 with DLBCL, 3 with transformed follicular lymphoma (tFL), and 1 with primary mediastinal B-cell lymphoma (PMBCL). Median age was 66 (range: 19-76), and 8 (29%) were female. The median number of prior treatments was 3 (range: 1-6), with 21 (75%) patients exposed to platinum-containing regimens and 5 patients (18%) had undergone high dose melphalan-based conditioning and autologous stem-cell rescue (HDT/ASCR). Nineteen patients (67.9%) received bridging therapy between apheresis and CAR-19 infusion. After treatment, 4 patients (14.3%) experienced grade 3 or higher cytokine release syndrome (CRS) and 9 patients (32.1%) had grade 3 or higher immune effector cell-associated neurotoxicity syndrome (ICANS). One patient passed away within a week post-infusion due to CAR-19 toxicity and with unknown disease response. This patient was omitted from progression free survival (PFS) but included in overall survival (OS) analyses. Median OS for the cohort as a whole was 11.6 months, with PFS 8.0 months (**Supplementary Figure 1a-b**). Durable responses were seen in 11 (39%), of which 10 were complete responses (CR) and one a durable partial response (PR) (**Table 1**). Overall, these results are comparable to previously reported axi-cel outcomes^4,9^.

**TABLE 1:** Patient Information.

| Characteristic | All Patients<br>(n = 28) |
| --- | --- |
| **Age, years** |  |
| Median | 66 |
| Range | 19 - 76 |
| **Sex, n (%)** |  |
| Female | 8 (28.6%) |
| Male | 20 (71.4%) |
| **Disease, n (%)** |  |
| DLBCL | 24 (85.7%) |
| TFL | 3 (10.7%) |
| PMBCL | 1 (3.6%) |
| **Stage at apheresis, n (%)** |  |
| I/II | 5 (17.9%) |
| III/IV | 23 (82.1%) |
| **IPI at apheresis, n (%)** |  |
| 1-2 | 7 (25.0%) |
| 3-5 | 21 (75.0%) |
| **ECOG at apheresis, n (%)** |  |
| 0-2 | 21 (75.0%) |
| 3-4 | 7 (25.0%) |
| **Prior treatment regimens, n** |  |
| Median | 3 |
| Range | 1-6 |
| **Salvage Chemotherapies** |  |
| Platinum compounds | 21 (75.0%) |
| Cisplatin | 5 (17.9%) |
| Carboplatin | 11 (39.3%) |
| Oxaliplatin | 5 (17.9%) |
| Melphalan | 5 (17.9%) |
| **Previous HDT/ASCR, n (%)** | 5 (17.9%) |
| **Bridging therapy, n (%)** |  |
| No | 8 (28.6%) |
| Yes | 19 (67.9%) |
| N/A | 1 (3.6%) |
| **Cytokine release syndrome (CRS), n (%)** |  |
| Grade 0 | 4 (14.3%) |
| Grade 1-2 | 20 (71.4%) |
| Grade 3-4 | 4 (14.3%) |
| **Immune effector cell-associated neurotoxicity syndrome (ICANS), n (%)** |  |
| Grade 0 | 7 (25.0%) |
| Grade 1-2 | 11 (39.3%) |
| Grade 3-4 | 9 (32.1%) |
| **Durable Response, n (%)** |  |
| CR (complete response) | 10 (35.7%) |
| PR (partial response) | 1 (3.6%) |
| SD (stable disease) | 0 (0%) |
| PD (progressive disease) | 16 (57.1%) |
| Unable to assess | 1 (3.6%) |

**Abbreviations:** DLBCL (diffuse large B cell lymphoma), TFL (transformed follicular lymphoma), PMBCL (primary mediastinal B cell lymphoma), IPI (international prognostic index), ECOG (Eastern cooperative oncology group), HDT/ASCR (high-dose therapy with autologous stem-cell rescue).

### Mutations in driver genes associated with CAR-19 outcome in r/r LBCL

We first examined markers associated with prognosis in previously untreated DLBCL, which in other series have not associated with CAR-19 outcome^5^. Double hit (DH), defined as cases with a chromosomal rearrangement in *MYC* together with rearrangement(s) in *BCL2* and/or *BCL6*, did not correlate with outcome (**Supplementary Figure 1c**). Nor did double-expression (DE) of MYC and BCL2 proteins by immunohistochemistry (IHC, **Supplementary Figure 1d**)^17–20^. In line with recent evidence^21,22^, high metabolic tumor volume (MTV) associated with inferior outcome in our cohort (p=0.019, **Supplementary Figure 1e**). Using the WGS data, we assigned all patients (28/28) to one of the genomic clusters described by Chapuy et al., predictive of outcome in newly diagnosed DLBCL^23^. Most cases fell into Cluster #2 or Cluster #3, which are characterized by *BCL2* alterations and mutations in chromatin modifiers like *KMT2D* and *CREBBP* (Cluster #2) or inactivation of *TP53* and recurrent chromosome-segment amplifications and deletions (Cluster #3). No patients fell into the more favorable Cluster #4, and only 2 were in Cluster #1, which also has a more favorable outcome, likely reflecting improved responses to prior treatments in these patients. Only one patient fell into Cluster #5, a subgroup with notoriously worse outcome, suggesting highly aggressive disease phenotype at relapse limits opportunities for CAR-19 referral (**Supplementary Figure 1f**). Notably, this Cluster #5 patient unfortunately rapidly progressed and passed away a few weeks after treatment. We also used the publicly available LymphGen classification algorithm^24^ and the majority of classified cases fell into the EZB cluster, characterized by epigenetic dysregulation and corresponding roughly to Cluster #3 in Chapuy (**Supplementary Figure 1g**)^24^. Neither system showed prognostic significance to CAR-19. In line with other reports, we found no established markers of prognosis in newly diagnosed DLBCL correlated with response to CAR-19, and we therefore initiated WGS-based unbiased definition of the key genomic resistance drivers.

Including the two cases with both pre- and post-CAR-19 samples, 30 total tumor samples successfully underwent WGS from the 28 r/r patients, together with matched germline for all individuals. We found a median of 12801.5 somatic variants per sample (range: 5382-28033 somatic variants) (**Supplementary Figure 2a**). Patients who progressed on CAR-19 had an increased number of variants compared to those who achieved a prolonged remission (p=0.034, **Supplementary Figure 2b**); however, there was no difference in total nonsynonymous mutational burden between patients who progressed on CAR-19 versus patients with durable responses. To identify driver genes, we leveraged the ratio of nonsynonymous to synonymous mutations using the dNdScv algorithm^25^. To increase statistical power, we combined our r/r cohort with 50 newly diagnosed DLBCL cases from the Pan-Cancer Analysis of Whole Genomes (PCAWG)^26^. Positive selection was detected in 36 candidate driver genes (q value < 0.1; **Supplementary Table 2**)^25^. After correction for multiple testing using false discovery rate (fdr), we found that only *TP53* was significantly enriched in our cohort (fdr=0.069) in comparison with the PCAWG cohort. It was also the most frequently mutated gene with 50% of r/r cases containing at least one mutation (**Supplemental Figure 2c**). Nevertheless, *TP53* did not predict poor CAR-19 outcome. These findings are consistent with what has been previously reported in r/r cases^27^. Among these positively selected driver genes and genes known to be involved in DLBCL pathogenesis (**Supplementary Figure 2c**), only NF-kappa-B-inhibitor-alpha (*NFKBIA)* and *MYC* mutations were associated with worse PFS after CAR-19 (p=0.04, p=0.025 respectively, **Supplementary Figure 2d-e**).

### Mutational signatures’ impact on CAR-19 efficacy

Next, we ran *sigProfiler*^28^ and hierarchical Dirichlet^29^ to investigate underlying mutational processes (i.e., mutational signatures) involved in shaping the repertoire of single base substitutions (SBS). To increase resolution and statistical power we combined our r/r cohort with 50 WGS from patients with newly diagnosed DLBCL included in PCAWG^26^. Combining these two de novo mutational signatures approaches, we identified 12 mutational signatures involved in our cohort of r/r lymphomas. Eight of these are currently included in the COSMIC catalog v.2 (https://cancer.sanger.ac.uk/signatures/sbs) and have previously been reported in newly diagnosed DLBCL: SBS1 (aging), SBS2 (APOBEC), SBS5 (aging), SBS8, SBS9 (poly eta - germinal center), SBS13 (APOBEC), SBS17b and SBS18 (reactive oxygen species)^28^. All of the other four extracted mutational signatures are caused by exposure to distinct chemotherapies and 2 are not yet included in the COSMIC catalog (SBS-MM1 = melphalan; E_SBS37 = oxaliplatin, SBS31 = cisplatin/carboplatin, SBS35 = cisplatin signatures; **Figure 1a**)^28–33^. Next, to confirm the presence of each mutational signature and to accurately estimate its contribution, we ran the *mmsig* fitting algorithm^34^. As expected SBS-MM1 was identified in 4 out of 5 patients who received melphalan as part of HDT/ASCR (**Figure 1b**). To accurately define evidence of platinum mutagenic activity, we implemented the double base substitution analysis (DBS) and detected DBS5 (platinum chemotherapy treatment signature) in 83% (15/18) of patients who had evidence of platinum SBS-signatures (**Figure 1b**). Interestingly 6 out of 24 (25%) previously exposed to platinum did not show any sign of these chemotherapy related SBS and DBS signatures. It has been shown that distinct chemotherapy agents promote their mutagenic activity introducing a unique catalogue of mutations in each exposed single cell^29,32,33,35^. Therefore, this single-cell chemotherapy-barcode will be detectable by bulk WGS only if one single tumor cell exposed to the chemotherapy expands, taking clonal dominance (i.e., single-cell expansion model). In contrast, chemotherapy-induced mutational signatures will not be detectable if the cancer progression is driven by multiple clones originating from different single cells exposed to chemotherapy and therefore harboring different chemotherapy-barcodes (catalogue of unique chemotherapy-related mutations). The concept of chemotherapy-barcoding can also be used to time events and to establish if the progression was driven by one or more single tumor cells^33^. To do so, we reconstructed the phylogeny of two cases with samples collected before and at relapse after CAR-19 therapy (**Methods**). In one patient (CAR_84), the clonal composition did not change over time and no platinum-related signatures were detected despite prior exposure, suggesting a complete refractoriness to CAR-T where the progression is driven by multiple tumor cells/clones. The other case (CAR_39) is an example of branching evolution after CAR-19, with each branch characterized by a unique SBS-MM1 catalogue of mutations (**Figure 1c**). This scenario is compatible with progression driven by a single cell previously exposed to melphalan (SBS-MM1) and platinum (SBS31) that took clonal dominance to drive relapse after initial complete remission in response to CAR-T infusion. Overall, these data revealed that, similar to other cancers^33,35^, aggressive lymphomas can increase mutational burden at relapse due to exposure to mutagenic agents, and progression can be driven by single surviving cells.

**Fig.1.**
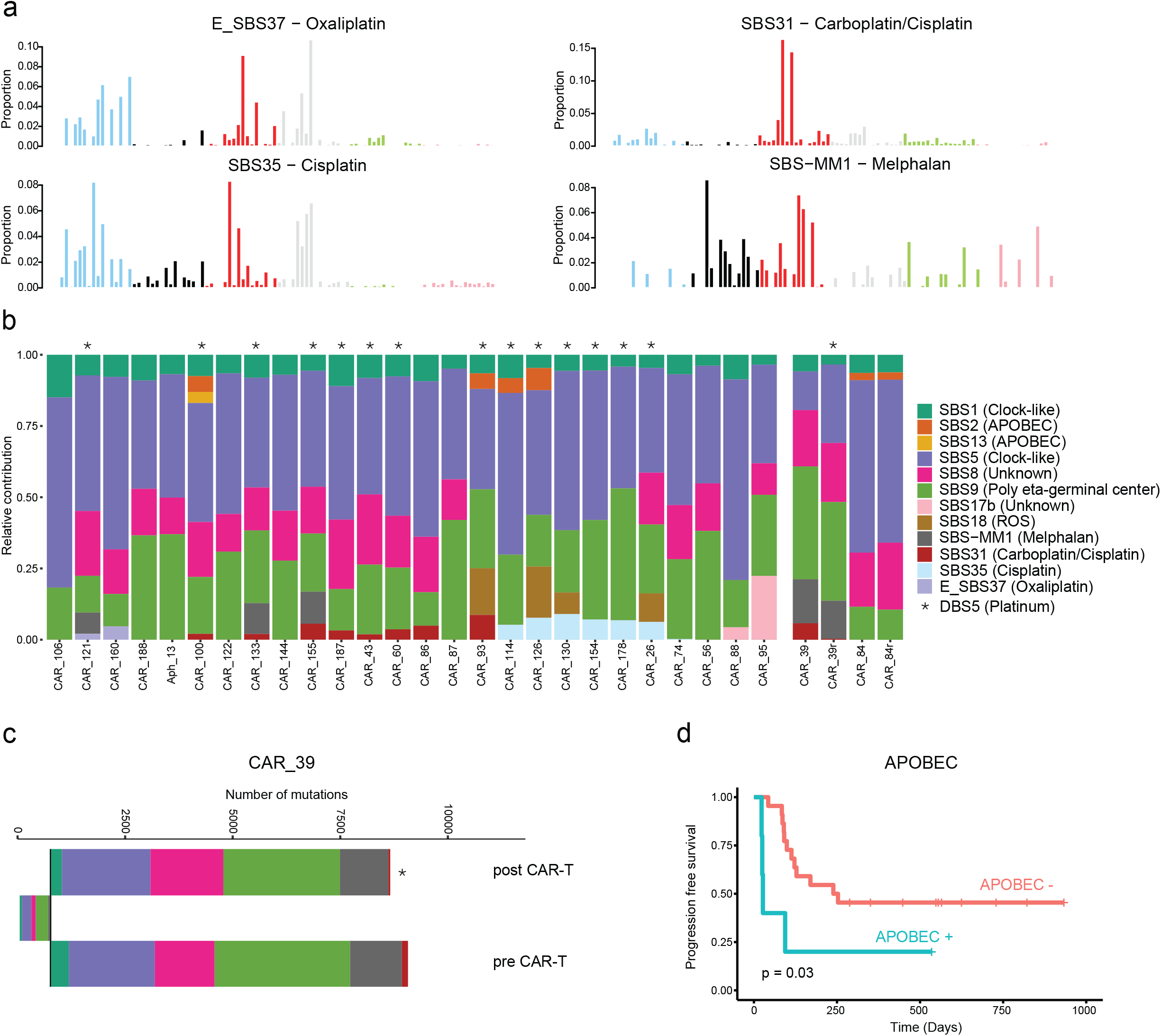
Relapsed or refractory large B-cell lymphoma mutational signatures landscape. **a)** The 4 chemotherapy-related signatures detected in our 28 r/r LBCL patients treated with CAR-T cell therapy. **b)** The relative contribution of each mutational signature (color) per each sample (x-ax-is). Asterisks indicate the presence of DBS5 (Platinum chemotherapy treatment double base substitution signature). **c)** Mutational-signature contributions for each phylogenetic tree cluster (sample CAR_39). Mutational signature colors are the same as the figure b legend. Asterisk indicates the present of DBS5. **d)** The Kaplan-Meier plot of progression free survival (PFS) comparing patients with (APOBEC +; in blue) and without APOBEC signature (APOBEC -; in red).

When correlated to response to CAR-T therapy, SBS2 and SBS13 (APOBEC) carried a significantly worse PFS with 4/5 patients progressing within four months (p=0.03; **Figure 1d**). APOBEC refers to a family of cytidine deaminases that generates an innate immune response to viruses and which has been shown to be active in many human cancers^28,29,36^, in particular in refractory tumors^37^, in metastasis^38,39^, and in tumors with loss of *HLA*^40^. Specific to lymphoma, APOBEC3 family members have been shown to contribute to lymphomagenesis in primary effusion lymphoma, and its mutagenic activity can be detected in 7.8% of newly diagnosed DLBCL^28,41^. APOBEC signatures in LBCL tumors may therefore be a biomarker of poor response to CAR-19.

### Focal deletions of *RB1* or *RHOA* and poor CAR-19 responses

We ran the *GISTIC v2*.*0* algorithm^42^ to compare the genome-wide CNV distribution between our 28 r/r patients and 50 newly diagnosed DLBCL in PCAWG^26^ (see **Methods**). We detected 8 arm-level and 8 focal regions of copy number gain and 3 arm-level and 19 focal regions of copy number loss (q value < 0.1; **Figure 2a** and **Supplementary Figure 3a**). Comparing the prevalence of these significant CNVs between r/r and *de novo* DLBCL three deletions emerged as statistically significant and enriched in the first group: chr17p13.1 (*TP53*; p=0.038), chr3p21.31 (*RHOA*; p=0.05), and chr13q14.2 (*RB1*; p=0.038; **Figure 2b**). Despite the high prevalence of *TP53* deletions in our r/r cohort (46.4%), this lesion did not carry prognostic impact in patients after CAR-19 (**Figure 2c**). Combining *TP53* with the related tumor suppressor *CDKN2A*, 78.6% of our samples had at least one mutated or deleted allele in one of the two genes (**Supplementary Figure 3b**), reflecting the aggressive nature of the tumors included in our cohort which had relapsed after multiple courses of intensive chemotherapy. Interestingly, deletions involving *RHOA* and *RB1* were strongly predictive of poor outcome after CAR-19 (p=0.0007 and p=0.05 respectively; **Figure 2c**) with 5/5 (100%) and 6/8 (75%) patients respectively whose tumors harbored these deletions progressing. This analysis highlights two novel driver genes in CAR-19 response in r/r LBCL and further confirms CAR-19 outcomes are affected by genomic features different from those associated with poor prognosis in newly diagnosed DLBCL.

**Fig.2.**
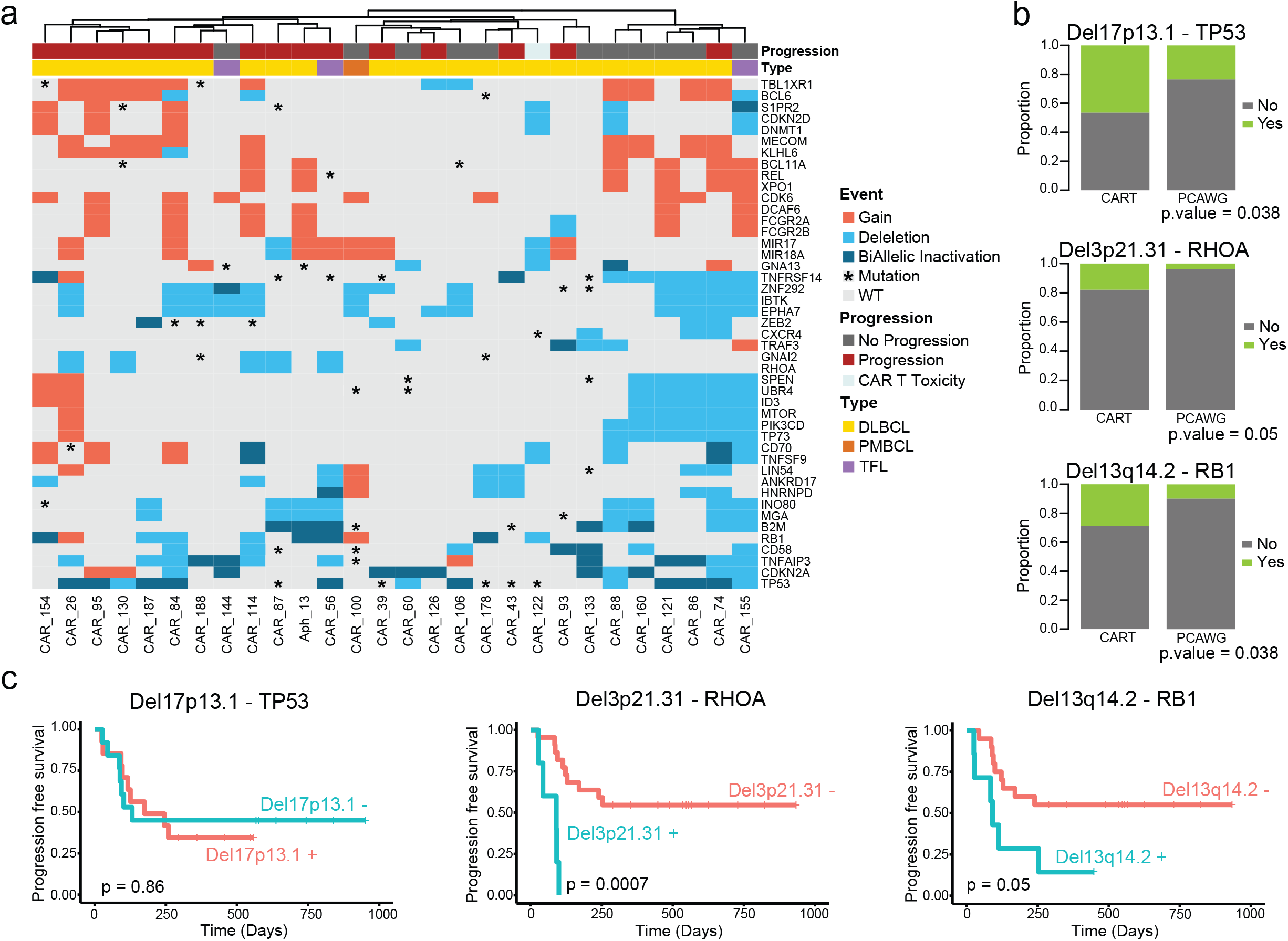
Clinical impact of recurrent copy number anomalies in r/r LBCL. **a)** The heatmap shows the significant genes extracted by GISTIC combining r/r LBCL and newly diagnosed PCAWG samples. The unsupervised hierarchical clustering was performed using Euclidean distance and Ward’s linkage. Biallelic inactivation is defined as the presence of either two deletions or one deletion and one mutation in the same patients. **b)** Stacked bars show the significant GISTIC peaks enriched in our r/r LBCL compare to PCAWG. The y-axis specifies prportions for each sample. The p value was obtained with one-tailed Fisher’s exact test. **c)** Kaplan-Meier plots showing the impact of TP53, RHOA and RB1 deletion on progression free survival after CAR-19 of progression free survival.

### Chromothripsis events mark cases doomed to fail CAR-19 treatment

WGS allows detailed identification of structural variants (SVs) and complex events. We identified a total of 1669 SVs across the 30 WGS samples (median 42.5 per r/r patients, range 9-156; **Figure 3a**). Similar to other hematologic malignancies^43–46^, we observed evidence of three main complex SV events: chromothripsis, chromoplexy, and templated insertion. Chromoplexy is defined as a concatenation of structural variants leading to multiple simultaneous chromosomal losses. Templated insertions represent a concatenation of interchromosomal structural variants leading to a derivative chromosome where multiple focal gains involving oncogenes and regulatory regions are strung together and reinserted in the genome^43,45^. Chromoplexy and templated insertions were observed in 32.1% and 25% of patients respectively, and only templated insertions were enriched in the cohort compared to PCAWG (p = 0.029). Chromothripsis represents a catastrophic event in which one or more than one chromosome is shattered and aberrantly reassembled generating multiple aneuploidies (**Figure 3b**)^43,47^. This event was identified in 39.3% of r/r cases, slightly higher than in newly diagnosed DLBCL (24%, **Figure 3a**) though not significantly enriched. Interestingly, across all different SV and complex events, only chromothripsis had a significant impact toward worse PFS (p=0.041, **Figure 3c**) after CAR-19 treatment, with 9/11 (81%) cases experiencing early progression. Chromothripsis has often been associated with presence of APOBEC in other cancers^44,45,47^, therefore we investigated the relationship between these two genomic features across our cohort and their impact on outcome post CAR-19. Interestingly, only 1/5 (20%) cases with APOBEC had evidence of chromothripsis, suggesting an absence of a strong relationship between these two features. Notably, patients with either APOBEC or chromothripsis were characterized by a particularly unfavorable PFS (p=0.0057, **Figure 3d**).

**Fig.3.**
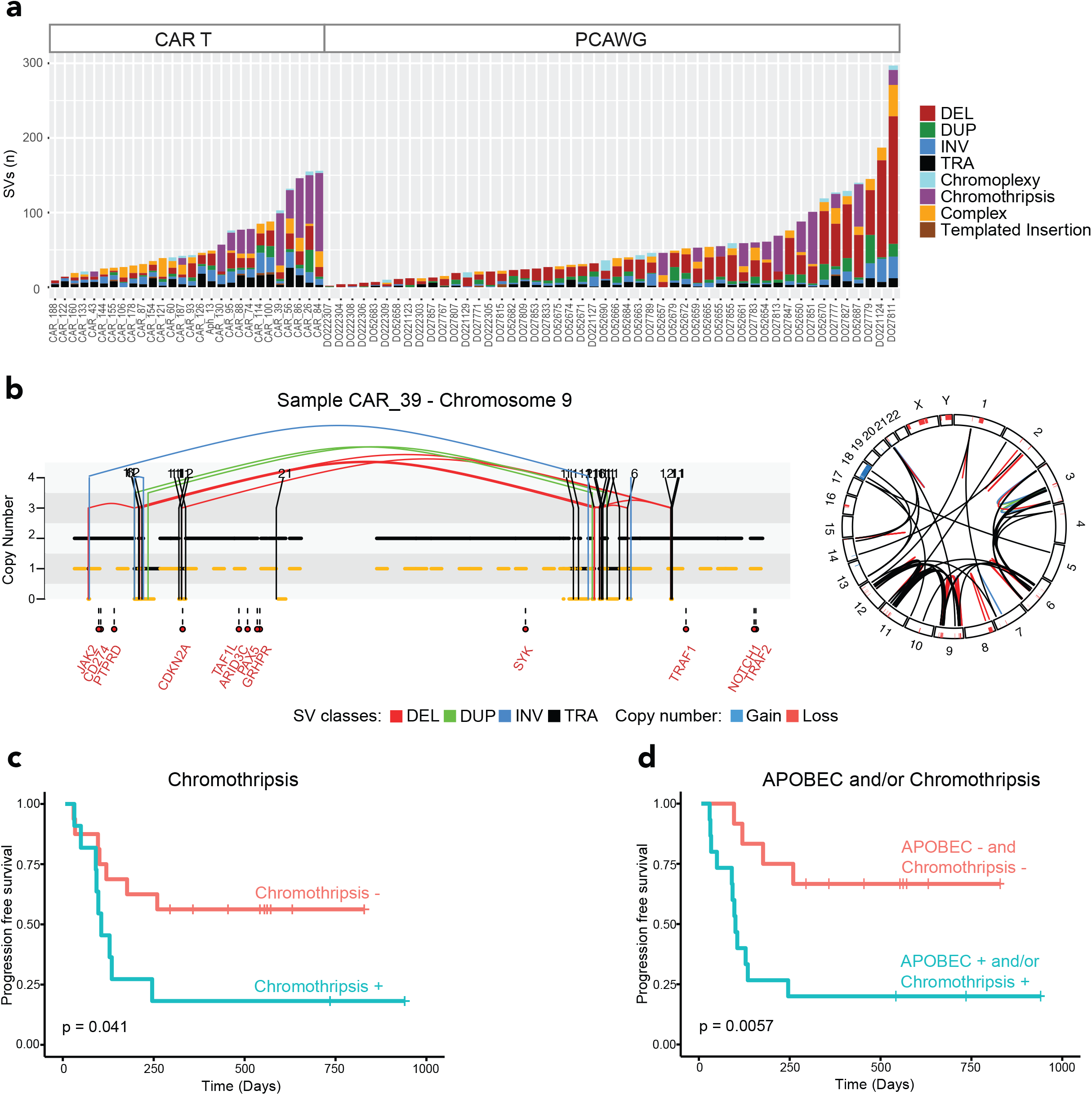
The landscape of structural variants in r/r LBCL and outcome association. **a)** Stacked bars show the genome-wide burden of each SV class and complex event per each sample (x-axis), grouped by analysis cohort. **b)** Left side, copy number profile plot integrated with SV information showing an emblematic example of chromothripsis on chromosome 9 responsible of *CDKN2A* loss (sample CAR_39). The horizontal black line indicates the total copy number; the dashed orange line indicates the minor copy number. The vertical lines represent SV breakpoints, color-coded based on SV class. Red text represents the DLBCL driver genes present on chromosome 9. Right side, the circos plot showing the genome wide distribution of the same chromothripsis event. **c)** Kaplan-Meier plot for progression free survival comparing patients with and without chromothripsis. **d)** Kaplan-Meier plot showing the poor progression free survival of patients with chromothripsis and/or APOBEC.

### Novel genomic features are detectable in nearly every CAR-19 failure

Here we identify a set of unique genomic features that correspond with poor prognosis for CAR-19 therapy in heavily pretreated, r/r LBCL patients. The most frequent and significant genomic features reported in this cohort and associated with poor outcome after CAR-19 treatment were: chromothripsis, *RB1* deletions, *RHOA* deletions, APOBEC mutational signature, *NFKBIA* mutations and *MYC* mutations (**Figure 4a** and **Supplementary Figure 5**). Of the patients that progressed, 15/16 (93%) had at least one of these genomics features and this translates into worse PFS (p=0.0028, **Figure 4b**). Individually, all genomic features correlated with significantly worse PFS but only the presence of *MYC* mutations, chromothripsis events and *RHOA* deletions correlated with significantly worse OS. (**Supplementary Figure 4**). Interestingly, these features do not overlap with previously reported negative prognostic indicators in DLBCL including rearrangements of *BCL2, BCL6*, or *MYC*^43–45^.

**Fig.4.**
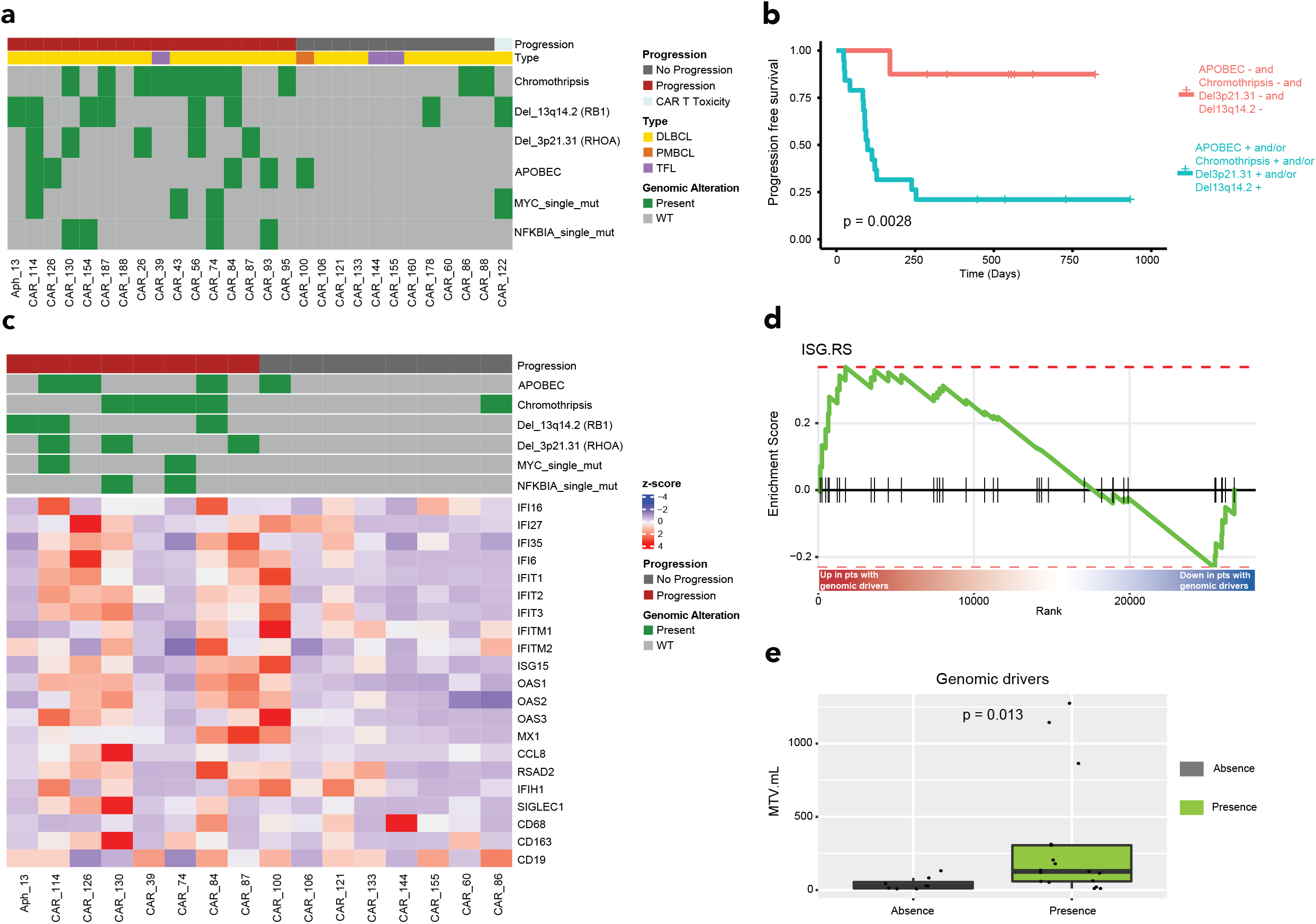
Impact of genomic alterations on clinical outcome and tumor microenviroment. **a)** The heat-map shows all the genomic alterations associated with progression after CAR-19 cell therapy. **b)** Prognostic impact of all the genomic alteration associated with progression after CAR-19 cell therapy. **c)** The heatmap shows the z scores for the IFN target genes, macrophage markers and CD19. **d)** Enrichment plot from GSEA showing the enrichment of the tumor IFN signature (ISG.RS) in patients with at least one significant genomic driver associated with progression after CAR-19 cell therapy. The vertical black bars indicate the position of the genes in the signature along the ranked gene list, the green line shows the enrichment score along the ranked gene list. The red to blue color bar shows the ranking of genes from up- to downregulated in patients with at least one genomic driver. **e)** The boxplot indicates the association between the metabolic tumor value (MTV in mL) and patients with at least one significant genomic driver. The p value was obtained with two tiled Fisher’s exact test.

It has been shown that distinct tumor immune microenvironmental patterns correlate with clinical outcome in patients treated with CAR-19^12,13^. In this study, we showed how distinct and complex genomic features in the tumor cells are strongly predictive of outcome in the same setting. To investigate a link between these two different assessments, we interrogated by RNA-seq the T-cell exhaustion landscape and IFN-signaling across 16 patients included in our study. The differential expression analysis (16 patients with also WGS data available) showed a higher expression level of genes known to be target of tumor signaling in patients treated with CAR-19 (**Figure 4c**). Interestingly, the ISG.RS signature, previously described to be associated with T-cell exhaustion and worse outcome after immunotherapy^12,48^, was enriched in r/r patients harboring at least one reported genomic driver (**Figure 4d**), while the INFG.GS signature, associated with higher response to check point blockade, was enriched in patients without any significant genomic drivers (**Supplementary Figure 6**). Moreover, the cases containing at least one reported genomic driver were characterized by higher MTV, reflecting the relationship between disease aggressiveness and the TME (**Figure 4e**). Overall, these data suggest resistance to CAR-19 in r/r aggressive lymphoma is mediated by a complex interplay between distinct tumor genomic and immune microenvironment features.

### Target-independent CAR-T anti-tumor activity

Past studies have assessed loss of CD19 as a mechanism of resistance to CAR-19 therapy^49^. Although individual examples of such cases demonstrate lack of response^50^, this mechanism of escape seems to explain only a small proportion of resistance in DLBCL^51^. In our cohort, pre-treatment CD19 expression was tested in 20 cases by flow cytometry, with 3 (15%) having reduced expression. Of these, two patients progressed, and one achieved a durable response. Additionally, all four CAR-19 relapse tumors were positive for CD19. At the genomic level, two cases showed a monoallelic copy number loss of *CD19* and one case had *CD19* subclonal mutation (L174V; 30% CCF). Strikingly, all three of these cases had durable CAR-19 responses. The last case with CD19 L174V is an emblematic example reflecting the complexity of the anti-tumor activity promoted by CAR-19. An identical mutation was found as clonal at baseline in a patient that was completely refractory to CAR-19 treatment (**Figure 5A, top**)^50^. In line with this prior evidence, in our patient we would have expected a CAR CD19 mediated eradication of all CD19 wild type (wt) cells, but not of the one harboring CD19 L174V mutation (**Figure 5a, middle**). However, the CAR-19 infusion induced a complete tumor eradication (i.e., both CD19 wt and CD19 L174V mutated clones) and an ongoing remission of more than 2 years post-CAR-19 infusion (**Figure 5a, bottom**). Taken together with 6/8 patients lacking CD19 who responded to axi-cel in the ZUMA-1 registration study^6^ and with recent pre-clinical studies^52,53^, these findings indicate antigen-mediated tumor killing is not the only mechanism of tumor eradication, and alternate mechanisms may predominate.

**Fig.5.**
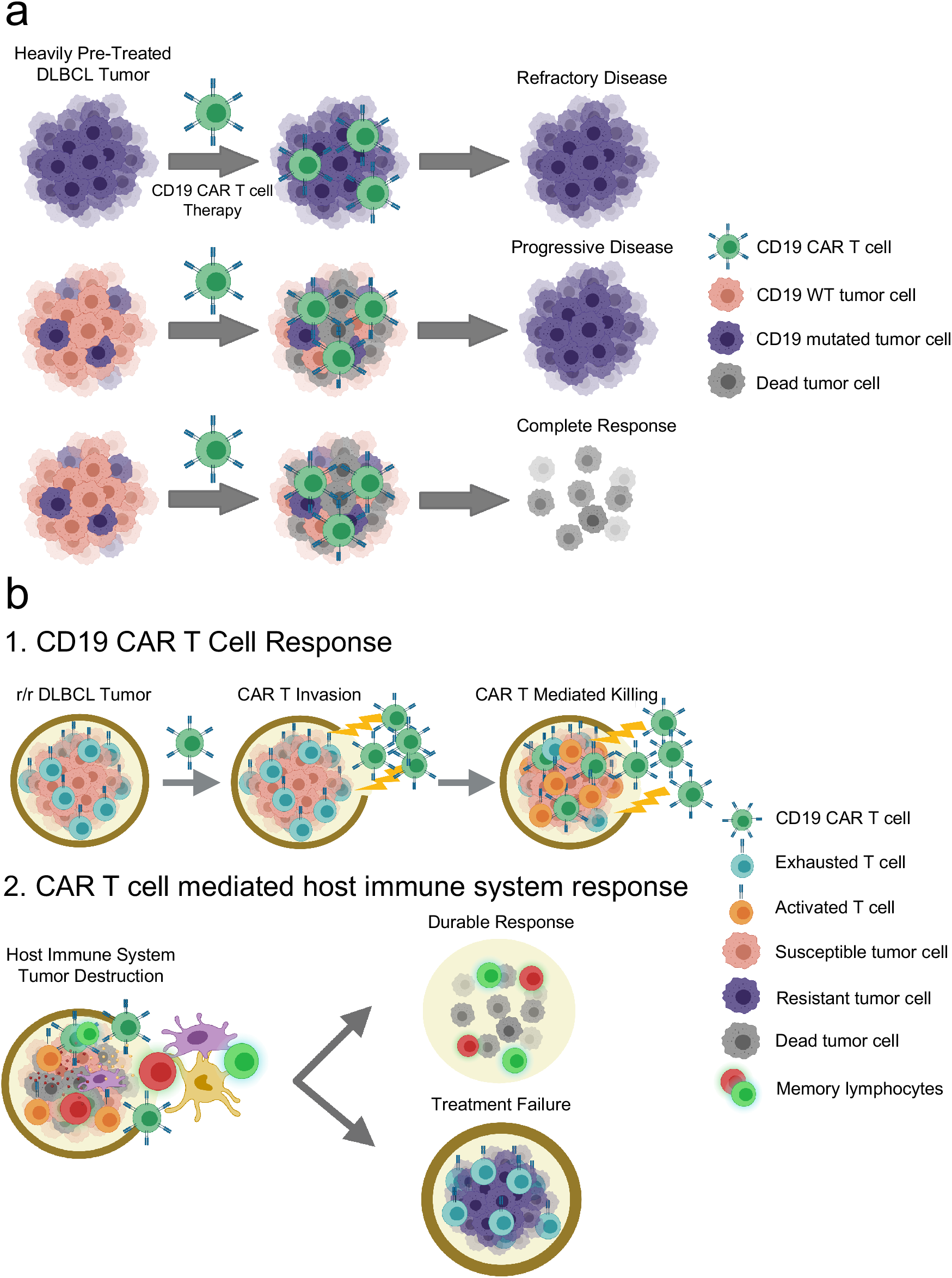
Model of the antigen-independent mechanisms of CAR-T mediated tumor-killing. **a)** Genetic alterations of *CD19* in tumor cells do not always affect CAR-19 outcomes in r/r LBCL. **b)** The anti-tumor CAR-19 activity can be summarized in two main phases: 1) CAR-19 cells invade the TME and initiate attack on tumor; 2) the subsequent full clearance depends on successful overall host response (2).

## DISCUSSION

To our knowledge, these data provide the first unbiased genome-wide discovery of tumor-intrinsic factors associated with resistance to CAR-T therapy. Genomic complexity indicated by evidence of chromothripsis events and APOBEC mutational activity was detected in most r/r lymphomas that progressed after CAR-T therapy. Strikingly and independently, focal deletions in *RB1* and *RHOA* also strongly correlated with a poor CAR-19 response in these heavily pretreated, r/r patients. Together, at least one of these findings was present in 15/16 (93.8%) of cases assessable for response and that progressed after therapy. These specific genomic findings not only provide biomarkers predictive of poor response to CAR-19 in r/r DLBCL patients but more importantly emphasize the need for functional studies to elucidate the mechanisms of these events in both primary and r/r disease. Some of these findings, such as chromothripsis and APOBEC, have been linked to more aggressive and resistant tumors^45,47^. The role of chromothripsis and APOBEC in newly diagnosed DLBCL has not been tested in prior studies, and the low number of progressed cases in the PCAWG dataset does not allow for a proper investigation (n=3). However, our data suggest patients with chromothripsis and/or APOBEC generally fail CAR-19 and present high-risk and exhausted microenvironmental patterns. The idea that complex genomic features and genomic instability are linked to a more immunosuppressed environment is not new^54^, but this is the first evidence from clinical samples showing it in the context of CAR-19 therapy.

Focal deletions of *RB1* or *RHOA* also correlated with lack of durable response to CAR-19. Given the crucial role of *RB1* in regulating cell cycle progression, the mechanism of progression in these patients could be directly related to overall tumor burden as the CAR-T response seems to not be able to overcome significant bulky disease^11,21^. As a matter of the fact, patients with *RB1* deletion had significant higher MTV compared to patients without any genomic features associated with CAR-19 resistance (p=0.026). The RHOA protein, meanwhile, affects a wide range of cellular processes in diffuse cell types, and its mechanisms as a DLBCL tumor suppressor remain to be clearly defined. Studies to date show increased motility of malignant and pre-malignant B lymphocytes in RHOA loss-of-function experiments^55,56^. Therefore, dissemination to tissues or niches in the TME that provide sanctuary from CAR-19-initiated immunity is one hypothesis. Given the prevalence of *RHOA* deletions in de novo DLBCL^23,57^ and now also correlation with poor prognosis to CAR-19, detailed laboratory studies are warranted to explore the role of these focal deletions in DLBCL and assess their impact on CAR-T response.

We provide cell-intrinsic alternatives to loss of the CD19 CAR target, a mechanism that, while logical, appears to be of unclear real-world importance^52,53^. In this cohort, genomic alterations of *CD19* or reduced expression by flow cytometry did not significantly affect outcome, revealing for the first time that the CD19-independent genomic drivers of CAR-19 resistance appear by far the more clinically important and inter-connected with the TME. Moreover, durable responses in cases with *CD19* monoallelic deletion or sub-clonal mutation demonstrate antigen-independent clearance can play a key role in clinical responses to CAR-19. Taken alone, these findings would be hypothesis generating, but they are in fact highly consistent with multiple published clinical and preclinical observations. For example, multiplex immunostaining of samples from axi-cel treated patients recently showed ≤5% of TME T cells were CAR-positive five days after infusion, but the CAR-negative cells were diffusely activated and likely contributed to both therapeutic efficacy and CRS toxicity^58^. Therefore, though individual case studies have implicated antigen loss as a mechanism of CAR-T resistance^49,50,59^, this does not account for the majority of resistant cases. Quantitative flow cytometry recently suggested lower pretreatment density of CD19 molecules per tumor cell associated with worse CAR-19 responses in LBCL^15^, but it was not possible to carry out this specialized assessment in our cohort for comparison to genotypes. We propose that an essential mechanism of CD19 CAR-T cells is to penetrate the exhausted TME providing access for the host immune system to attack the tumor (**Figure 5b**). In this scenario, the CAR-T cells act as a gateway into the immunosuppressed TME to allow the host immune system to destroy the tumor. Data from axi-cel-treated patients showed that patients with high serum inflammatory markers, along with increased tumor IFN signaling were indicative of lack of durable response^12^. Recent studies by Alizadeh et al^53^ demonstrated that CAR-T cells secrete IFN gamma and activate host T cells in a mouse model of glioblastoma and these cells preserved their anti-tumor activity also when infused without CAR-T. Combining these results with our genomics data, the role of the CAR-T cells in invigorating the host immune response in an exhausted TME emerges as the key to maintaining a durable response to CAR-T in r/r DLBCL patients. At the same time our data reveal that genomically complex and unstable tumors have a high degree of exhaustion and immunosuppression, and this creates a perfect storm of conditions for limiting the CAR-T activity and clinical efficacy.

Many CAR-T products are under development for use in various solid and hematologic malignancies with mixed efficacies^1,2^. This model of CAR-T resistance might be applicable to diseases such as multiple myeloma, where, APOBEC and chromothripsis are more frequent and an even higher percentage of patients relapse after CAR-T compared to DLBCL^29,45,46,60^. Clearly, further research is warranted to understand the role of the CAR-19 cells on the tumor microenvironment and subsequently identify ways to bolster the response to CAR-T through reactivation of the host immune system against the tumor.

## METHODS

### Patients and Samples

#### Patients

Patient characteristics and clinical outcomes for our cohort of 31 r/r LBCL patients are recorded in Table 1 and germline and tumor samples were collected prospectively following established international review board protocols^12^. Research was conducted in accordance with the Declaration of Helsinki. WGS was performed for patients with adequate samples at the time of analysis without further selection. Durable responders (non-progressors) were defined as patients who maintained remission after a minimum follow-up of 6 months after CAR19 infusion. Non-durable responders (progressors) had lymphoma recurrence or died from any cause.

To increase the statistical power in several analyses we included in this study 50 newly diagnosed DLBCL cases from PCAWG^26^, after removing the sample carrying the BRCA mutation.

#### Sample Collection & DNA Extraction

Patient samples were received as frozen peripheral blood mononuclear cells or viably preserved tumor biopsies and then thawed at 37C in a water bath. Once thawed, the samples were spun at room temperature at 3000g for 5 minutes. The cell pellets were washed once with phosphate buffered saline before being processing for nucleic acid extraction. The AllPrep DNA/RNA Mini Kit (Qiagen, Cat. #80284) was used to extract DNA and the samples were eluted in water.

#### Whole Genome Sequencing

WGS library construction and sequencing were performed at the Center for Genome Technology at the John P. Hussman Institute for Human Genomics, University of Miami Miller School of Medicine. First, all DNA samples were evaluated for concentration by fluorometric Qubit assays (Thermo-Fisher) and for integrity by TapeStation (Agilent Technologies). Sequencing libraries were prepared using the TruSeq DNA PCR-free HT sample preparation kit from Illumina. Briefly, one ug of total genomic DNA was fragmented using the Covaris LE220 focus acoustic sonicator to a target size of 350bp. Blunt-end DNA fragments were generated and size selection performed with AMPure bead purification (Beckman Coulter). A-base tailing was performed on the 3’ blunt ends followed by adapter ligation and a bead-based clean-up of the libraries. Final library fragment size was evaluated on the TapeStation (Agilent Technologies) and final molarity quantification determined by qPCR with adapter specific primers (Kapa Biosystems) on a Roche Light Cycler. Libraries were normalized to 2.8nM and 24-samples pooled for sequencing on a S4-300 flow cell on the NovaSeq 6000. Paired-end 150bp reads were generated to yield an average depth of 30x per sample. FASTQ files were generated using the Illumina BCL2FASTQ algorithm and used for downstream processing.

#### Whole genome sequencing analysis

Raw FASTQ files were uploaded to the Illumina BaseSpace Sequence Hub for downstream processing. Tumor and normal paired samples were aligned against the GRCh38 genome build and somatic single nucleotide variants (SNVs) and short insertion-deletions variants (indels) were called using the DRAGEN Somatic Pipeline Version 3.6.3. We performed additional filters to the only “PASS” calls, to remove artifactual variants. We excluded variants based on at least 1 of following filters: calls were unidirectional; an alternative allele was present in matched normal; the C>A/G>T variants had a frequency <0.1 (oxoG artifacts). We applied the *dN/dScv* method to detect genes under positive selection in r/r cases. To increase the statistical power we included 50 newly diagnosed DLBCL samples from PCAWG^26^. The algorithm estimates the excess of nonsynonymous mutations while accounting for the mutational spectrum and gene-specific mutation rates^25^. Then, we evaluated which results were enriched in our cohort using a two-tailed Fisher’s exact test and correction for multiple testing using false discovery rate (FDR).

CNVs were called with Sequenza Version 3.0.0 algorithm^61^ as previously described in DLBCL^62^. The genome regions that were significantly modified in our samples were identified by using *GISTIC* (v2.0.23)^42^. To improve the test’s statistical power, we run our r/r samples (n = 28) with the baseline DLBCL PCAWG samples (n = 50). In this way we were able to detect the anomalous peaks shared among all the samples, and subsequently to identify which of these were enriched in our r/r cohort, using a one tailed Fisher’s exact test. The analysis was executed using Gene Pattern web interface (http://genepattern.broadinstitute.org) and setting a q value threshold of 0.01. To determinate the tumor clonal architecture, and to model clusters of clonal and sub-clonal points mutations, we combined SNV and CNV data using the *PyClone-VI* (v0.1.0)^63^.

Mutational signature analysis of SBS was performed following three main steps: 1) *de novo* extraction, 2) assignment, and 3) fitting^31^. For the novo extraction of mutational signatures, we run *SigProfiler* and hdp algorithms^28,29^, combining our 28 r/r samples together with 50 baseline PCAWG samples. Next, the extracted process active in our cohort was assigned to one or more mutational signatures included in the latest COSMIC v3.2 catalog (https://cancer.sanger.ac.uk/signatures/sbs). Finally, the 28 r/r samples were run with a fitting algorithm designed for hematological cancers, *mmsig*^34^. It confirms and estimates the contribution of each mutational signature in each sample. Confidence intervals were generated by drawing 1000 mutational profiles from the multinomial distribution, each time repeating the signature fitting procedure, and finally taking the 2.5^th^ and 97.5^th^ percentile for each signature. Mutational signature analysis of DBS was performing using *SigProfiler* algorithm to *de novo* extraction and assignment with COSMICv.2 signatures catalog, combining our cohort with PCAWG cohort.

To detect the SVs, deletions, inversion, translocations and tandem duplications, we used *Manta*. Complex events such as chromothripsis, chromoplexy, templated insertions were defined after manual inspection as previously described^43,45,46,64,65^. Templated insertions were defined by translocations associated with copy number gain, resulting in concatenation of amplified segments from two or more chromosomes into a continuous stretch of DNA, inserted into any of the involved chromosomes. Chromoplexy connected segments from multiple chromosomes, but they are associated with copy number loss. Chromothripsis was defined by the presence of 10 or more interconnected SV breakpoint pairs associated with a shattering and random rejoining of one or more chromosomes with oscillating copy number^64^. Patterns of three or more interconnected breakpoint pairs that did not fall into either of the above categories were classified as unspecified complex. All SVs not part of a complex event were classified as single^45,46^.

#### RNA sequencing analysis

The RNA sequencing (RNA-seq) libraries were prepared with the NuGen RNA-Seq Multiplex System (Tecan US) as previously described^12^. The libraries were sequenced on the Illumina NextSeq 500 system with a 75 base paired end run at 80 to 100 million read pairs per sample. RNA-seq reads were mapped to the reference human genome (GRCh38) using the STAR algorithm^66^, and the parameters were set to count read numbers per gene while mapping. To analyze the gene expression profile, we used the DESeq2 R package^67^. First, the dataset of raw counts was filtered to remove genes with <10 reads in >95% of samples. Then, we performed the library size normalization, followed by the gene expression analysis. Nominal p values were corrected for multiple testing by using the Benjamini-Hochberg FDR method.

The Gene Set Enrichment Analysis (GSEA)^68^ was performed using the fgsea^69^ R package. The H Hallmark gene sets collection, retrieved from MSigDb database v 7.4^70^, was enriched with two INF signatures, ISG.RS and IFNG.GS, previously described to be associated with response to immunotherapy^12,48^. Genes were ranked using the statistic derived from differential expression analysis with DESeq function.

#### Chapuy et al Clustering

Samples were clustered according to the Chapuy clustering system methods^23^ using the SV, CNV and mutation data.

#### LymphGen Clustering

Using the publicly available LymphGen Classifier^24^ (https://llmpp.nih.gov/lymphgen/index.php), samples were categorized into the various subtypes based on the SV, CNV and mutational data.

#### Statistics

The comparison tests have been performed with Fisher’s exact test. Association of categorial variables with progression free survival (PFS) and overall survival (OS) were performed in a univariable fashion using Kaplan-Meier curves and a log-rank test. All analyses were performed in R, the language and environment for statistical computing (R Core Team, 2021).

## DATA AVAILABILITY

Submission of raw data to the European Genome-phenome Archive (EGA) is in progress. PCAWG data are available at https://dcc.icgc.org/ and EGAS00001001692 [https://ega-archive.org/studies/EGAS00001001692];

## ACKNOWLEDGEMENTS

This work was supported by grant award from the Florida Academic Cancer Center Alliance (to J.H.S. and M.L.D.) and the Sylvester Comprehensive Cancer Center NCI Core Grant (P30 CA 240139 to J.H.S and F.M.).

F.M. is supported by the American Society of Hematology.

## AUTHOR CONTRIBUTIONS

F.M., F.L.L., J.H.S., M.L.D. designed and supervised the study, collected, and analyzed the data and wrote the paper.

C.A.C., B.Z., M.D.J., collected, analyzed and interpreted the data and wrote the paper.

A.J.G. analyzed the data

O.L., R.F., and K.M.R collected the data

All authors read, revised, proofed the manuscript

## REFERENCES

1. Brudno, J. N. & Kochenderfer, J. N. Chimeric antigen receptor T-cell therapies for lymphoma. Nat. Rev. Clin. Oncol. 15, 31–46 (2018).

2. Sterner, R. C. & Sterner, R. M. CAR-T cell therapy: current limitations and potential strategies. Blood Cancer J. 11, 69 (2021).

3. Vairy, S., Lopes Garcia, J., Teira, P. & Bittencourt, H. CTL019 (tisagenlecleucel): CAR-T therapy for relapsed and refractory B-cell acute lymphoblastic leukemia. Drug Des. Devel. Ther. Volume 12, 3885–3898 (2018).

4. Locke, F. L. et al. Phase 1 Results of ZUMA-1: A Multicenter Study of KTE-C19 Anti-CD19 CAR T Cell Therapy in Refractory Aggressive Lymphoma. Mol. Ther. 25, 285–295 (2017).

5. Locke, F. L. et al. Long-term safety and activity of axicabtagene ciloleucel in refractory large B-cell lymphoma (ZUMA-1): a single-arm, multicentre, phase 1–2 trial. Lancet Oncol. 20, 31–42 (2019).

6. Neelapu, S. S. et al. Axicabtagene Ciloleucel CAR T-Cell Therapy in Refractory Large B-Cell Lymphoma. N. Engl. J. Med. 377, 2531–2544 (2017).

7. Nastoupil, L. J. et al. Standard-of-Care Axicabtagene Ciloleucel for Relapsed or Refractory Large B-Cell Lymphoma: Results From the US Lymphoma CAR T Consortium. J. Clin. Oncol. 38, 3119–3128 (2020).

8. Schuster, S. J. et al. Chimeric Antigen Receptor T Cells in Refractory B-Cell Lymphomas. N. Engl. J. Med. 377, 2545–2554 (2017).

9. Schuster, S. J. et al. Tisagenlecleucel in Adult Relapsed or Refractory Diffuse Large B-Cell Lymphoma. N. Engl. J. Med. 380, 45–56 (2019).

10. Jacobson, C. A. et al. Axicabtagene Ciloleucel in the Non-Trial Setting: Outcomes and Correlates of Response, Resistance, and Toxicity. J. Clin. Oncol. JCO.19.02103 (2020) doi:10.1200/JCO.19.02103.

11. Vercellino, L. et al. Predictive factors of early progression after CAR T-cell therapy in relapsed/refractory diffuse large B-cell lymphoma. 4, 9 (2020).

12. Jain, M. D. et al. Tumor interferon signaling and suppressive myeloid cells associate with CAR T cell failure in large B cell lymphoma. Blood blood.2020007445 (2021) doi:10.1182/blood.2020007445.

13. Deng, Q. et al. Characteristics of anti-CD19 CAR T cell infusion products associated with efficacy and toxicity in patients with large B cell lymphomas. Nat. Med. 26, 1878–1887 (2020).

14. Wang, N. et al. Efficacy and safety of CAR19/22 T-cell cocktail therapy in patients with refractory/relapsed B-cell malignancies. Blood 135, 17–27 (2020).

15. Spiegel, J. Y. et al. CAR T cells with dual targeting of CD19 and CD22 in adult patients with recurrent or refractory B cell malignancies: a phase 1 trial. Nat. Med. 1–13 (2021) doi:10.1038/s41591-021-01436-0.

16. Shah, N. N. et al. Bispecific anti-CD20, anti-CD19 CAR T cells for relapsed B cell malignancies: a phase 1 dose escalation and expansion trial. Nat. Med. 26, 1569–1575 (2020).

17. Collinge, B. et al. The impact of MYC and BCL2 structural variants in tumors of DLBCL morphology and mechanisms of false-negative MYC IHC. Blood 137, 2196–2208 (2021).

18. Ennishi, D. et al. Double-Hit Gene Expression Signature Defines a Distinct Subgroup of Germinal Center B-Cell-Like Diffuse Large B-Cell Lymphoma. J. Clin. Oncol. JCO.18.01583 (2018) doi:10.1200/JCO.18.01583.

19. Green, T. M. et al. Immunohistochemical Double-Hit Score Is a Strong Predictor of Outcome in Patients With Diffuse Large B-Cell Lymphoma Treated With Rituximab Plus Cyclophosphamide, Doxorubicin, Vincristine, and Prednisone. J. Clin. Oncol. 30, 3460–3467 (2012).

20. Johnson, N. A. et al. Concurrent Expression of MYC and BCL2 in Diffuse Large B-Cell Lymphoma Treated With Rituximab Plus Cyclophosphamide, Doxorubicin, Vincristine, and Prednisone. J. Clin. Oncol. 30, 3452–3459 (2012).

21. Dean, E. A. et al. High metabolic tumor volume is associated with decreased efficacy of axicabtagene ciloleucel in large B-cell lymphoma. 4, 9 (2020).

22. Malek, E. et al. Metabolic tumor volume on interim PET is a better predictor of outcome in diffuse large B-cell lymphoma than semiquantitative methods. Blood Cancer J. 5, e326–e326 (2015).

23. Chapuy, B. et al. Molecular subtypes of diffuse large B cell lymphoma are associated with distinct pathogenic mechanisms and outcomes. Nat. Med. 24, 679–690 (2018).

24. Wright, G. W. et al. A Probabilistic Classification Tool for Genetic Subtypes of Diffuse Large B Cell Lymphoma with Therapeutic Implications. Cancer Cell 37, 551-568.e14 (2020).

25. Martincorena, I. et al. Universal Patterns of Selection in Cancer and Somatic Tissues. Cell 171, 1029-1041.e21 (2017).

26. The ICGC/TCGA Pan-Cancer Analysis of Whole Genomes Consortium. Pan-cancer analysis of whole genomes. Nature 578, 82–93 (2020).

27. Rushton, C. K. et al. Genetic and evolutionary patterns of treatment resistance in relapsed B-cell lymphoma. 4, 13 (2020).

28. Alexandrov, L. B. et al. The repertoire of mutational signatures in human cancer. Nature 578, 28 (2020).

29. Rustad, E. H. et al. Timing the initiation of multiple myeloma. Nat. Commun. 11, 1917 (2020).

30. Maura, F. et al. The mutagenic impact of melphalan in multiple myeloma. Leukemia (2021) doi:10.1038/s41375-021-01293-3.

31. Maura, F. et al. A practical guide for mutational signature analysis in hematological malignancies. Nat. Commun. 10, 12 (2019).

32. Pich, O. et al. The mutational footprints of cancer therapies. Nat. Genet. 51, 1732–1740 (2019).

33. Landau, H. J. Accelerated single cell seeding in relapsed multiple myeloma. Nat. Commun. 11, 10 (2020).

34. Rustad, E. H. et al. mmsig: a fitting approach to accurately identify somatic mutational signatures in hematological malignancies. Commun. Biol. 4, 424 (2021).

35. Kucab, J. E. et al. A Compendium of Mutational Signatures of Environmental Agents. Cell 177, 821-836.e16 (2019).

36. Petljak, M. & Maciejowski, J. Molecular origins of APOBEC-associated mutations in cancer. DNA Repair 94, 102905 (2020).

37. Maura, F. et al. Biological and prognostic impact of APOBEC-induced mutations in the spectrum of plasma cell dyscrasias and multiple myeloma cell lines. Leukemia 32, 1043–1047 (2018).

38. Dentro, S. C. et al. Characterizing genetic intra-tumor heterogeneity across 2,658 human cancer genomes. Cell 184, 2239-2254.e39 (2021).

39. Priestley, P. et al. Pan-cancer whole-genome analyses of metastatic solid tumours. Nature 575, 210–216 (2019).

40. McGranahan, N. et al. Allele-Specific HLA Loss and Immune Escape in Lung Cancer Evolution. Cell 171, 1259-1271.e11 (2017).

41. Wagener, R. et al. Analysis of mutational signatures in exomes from B-cell lymphoma cell lines suggest APOBEC3 family members to be involved in the pathogenesis of primary effusion lymphoma. Leukemia 4 (2015) doi:doi:10.1038/leu.2015.22.

42. Mermel, C. H. et al. GISTIC2.0 facilitates sensitive and confident localization of the targets of focal somatic copy-number alteration in human cancers. Genome Biol. 12, R41 (2011).

43. Li, Y. et al. Patterns of somatic structural variation in human cancer genomes. Nature 578, 112–121 (2020).

44. Hadi, K. et al. Distinct Classes of Complex Structural Variation Uncovered across Thousands of Cancer Genome Graphs. Cell 183, 197-210.e32 (2020).

45. Rustad, E. H. et al. Revealing the Impact of Structural Variants in Multiple Myeloma. Blood Cancer Discov. 1, 258–273 (2020).

46. Maura, F. et al. Genomic landscape and chronological reconstruction of driver events in multiple myeloma. Nat. Commun. 10, 3835 (2019).

47. Cortés-Ciriano, I. et al. Comprehensive analysis of chromothripsis in 2,658 human cancers using whole-genome sequencing. Nat. Genet. 52, 331–341 (2020).

48. Benci, J. L. et al. Tumor Interferon Signaling Regulates a Multigenic Resistance Program to Immune Checkpoint Blockade. Cell 167, 1540-1554.e12 (2016).

49. Majzner, R. G. & Mackall, C. L. Tumor Antigen Escape from CAR T-cell Therapy. Cancer Discov. 8, 1219–1226 (2018).

50. Zhang, Z. et al. Point mutation in CD19 facilitates immune escape of B cell lymphoma from CAR-T cell therapy. J Immunother Cancer 11 (2020) doi:10.1136/jitc-2020-001150.

51. Chong, E. A., Ruella, M. & Schuster, S. J. Five-Year Outcomes for Refractory B-Cell Lymphomas with CAR T-Cell Therapy. N. Engl. J. Med. 2 (2021).

52. Boulch, M. et al. A cross-talk between CAR T cell subsets and the tumor microenvironment is essential for sustained cytotoxic activity. Sci. Immunol. 6, 4344 (2021).

53. Alizadeh, D. et al. IFNg is Critical for CAR T Cell Mediated Myeloid Activation and Induction of Endogenous Immunity. Cancer Discov. 29 (2021) doi:10.1158/2159-8290.CD-20-1661.

54. Hegde, P. S. & Chen, D. S. Top 10 Challenges in Cancer Immunotherapy. Immunity 52, 19 (2020).

55. Jiang, X. et al. HGAL, a germinal center specific protein, decreases lymphoma cell motility by modulation of the RhoA signaling pathway. Blood 116, 5217–5227 (2010).

56. Muppidi, J. R. et al. Loss of signalling via Gα13 in germinal centre B-cell-derived lymphoma. Nature 516, 254–258 (2014).

57. O’Hayre, M. et al. Inactivating mutations in GNA13 and RHOA in Burkitt’s lymphoma and diffuse large B-cell lymphoma: a tumor suppressor function for the Gα 13 /RhoA axis in B cells. Oncogene 35, 3771–3780 (2016).

58. Chen, P.-H. et al. Activation of CAR and non-CAR T cells within the tumor microenvironment following CAR T cell therapy. JCI Insight 5, (2020).

59. Sotillo, E. et al. Convergence of Acquired Mutations and Alternative Splicing of CD19 Enables Resistance to CART-19 Immunotherapy. Cancer Discov. 5, 25 (2015).

60. Munshi, N. C. et al. Idecabtagene Vicleucel in Relapsed and Refractory Multiple Myeloma. N. Engl. J. Med. 384, 705–716 (2021).

61. Favero, F. et al. Sequenza: allele-specific copy number and mutation profiles from tumor sequencing data. Ann. Oncol. 26, 64–70 (2015).

62. Arthur, S. E. et al. Genome-wide discovery of somatic regulatory variants in diffuse large B-cell lymphoma. Nat. Commun. 9, 4001 (2018).

63. Gillis, S. & Roth, A. PyClone-VI: scalable inference of clonal population structures using whole genome data. BMC Bioinformatics 21, 571 (2020).

64. Maciejowski, J. et al. APOBEC3-dependent kataegis and TREX1-driven chromothripsis during telomere crisis. Nat. Genet. 52, 884–890 (2020).

65. Korbel, J. O. & Campbell, P. J. Criteria for Inference of Chromothripsis in Cancer Genomes. Cell 152, 1226–1236 (2013).

66. Dobin, A. et al. STAR: ultrafast universal RNA-seq aligner. Bioinformatics 29, 15–21 (2013).

67. Love, M. I., Huber, W. & Anders, S. Moderated estimation of fold change and dispersion for RNA-seq data with DESeq2. Genome Biol. 15, 550 (2014).

68. Subramanian, A. et al. Gene set enrichment analysis: a knowledge-based approach for interpreting genome-wide expression profiles. Proc. Natl. Acad. Sci. U. S. A. 102, 15545–15550 (2005).

69. Korotkevich, G., Sukhov, V. & Sergushichev, A. Fast gene set enrichment analysis. 060012 https://www.biorxiv.org/content/10.1101/060012v2 (2019) doi:10.1101/060012.

70. Liberzon, A. et al. Molecular signatures database (MSigDB) 3.0. Bioinformatics 27, 1739–1740 (2011).

