## Supplementary Figures and Tables for "Genomic drivers of large B-cell lymphoma resistance to CD19 CAR-T therapy"

**Supplementary Figure 1. Kaplan-Meier curves of progression free survival for whole cohort and clinical parameter.**

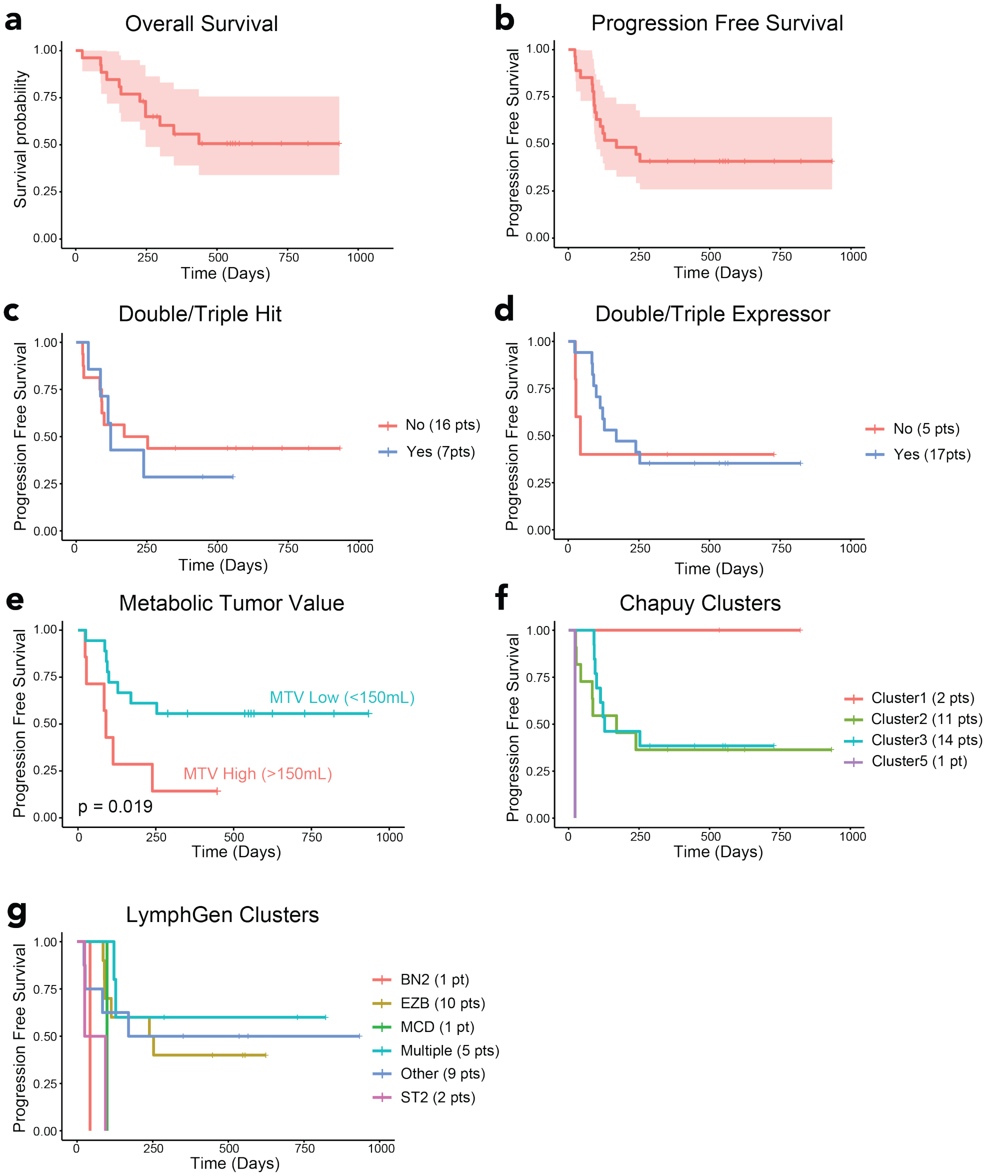

**Supplementary Figure 2. Mutational landscape in r/r DLBCL. a** Barplot shows the number of variants per each sample. **b** Boxplot representing the distribution of variant number in progression (blue) and no progression (yellow) samples. **c** The heatmap shows the DLBCL driver genes present in at least 10% of samples. Asterisks represent the significant gene in dN/dScv analysis (p < 0.1). **d-e** Kaplan-Meier curves of progression free survival.

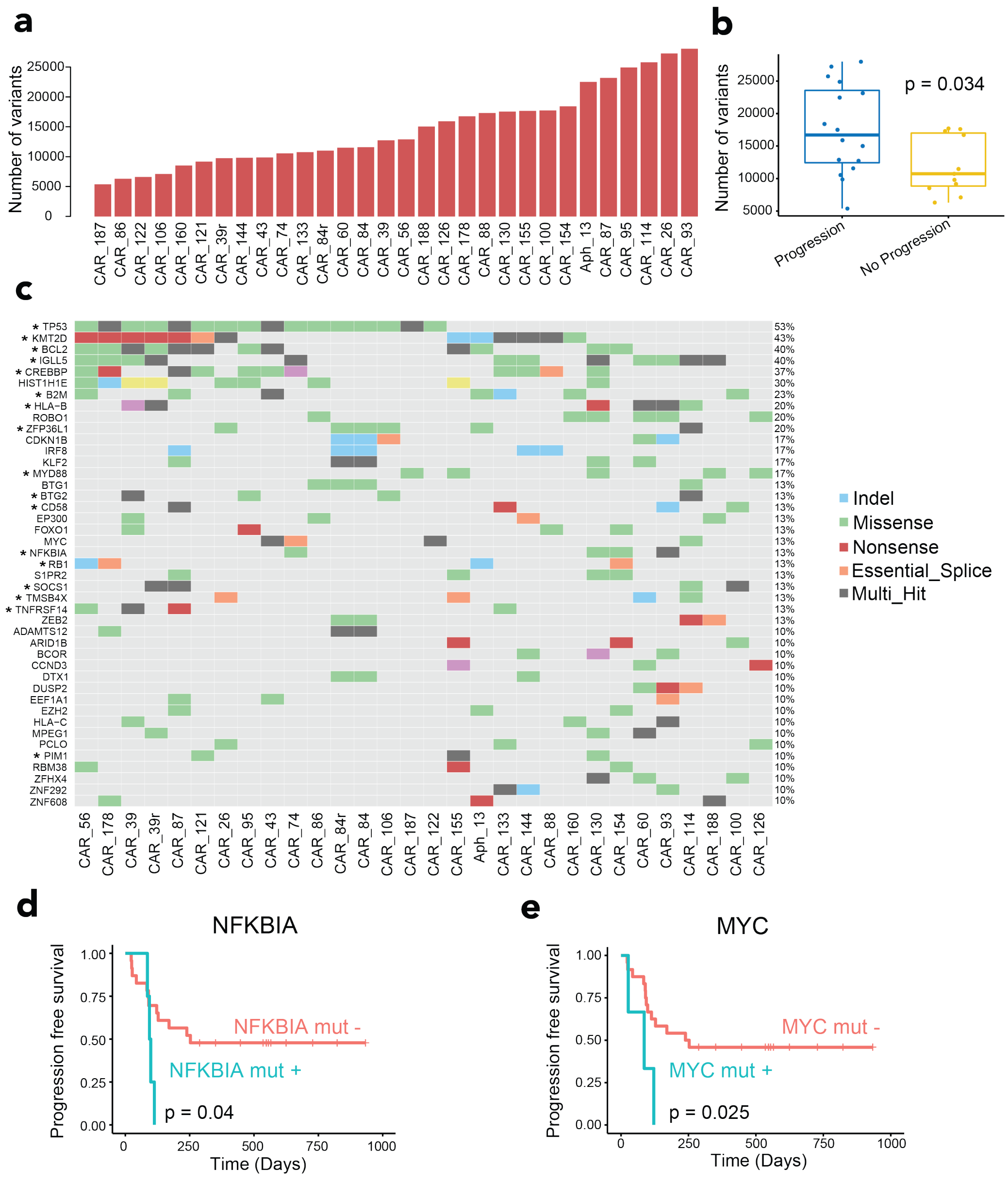

**Supplementary Figure 3. Recurrent copy number landscape in r/r DLBCL. a** Gistic significant peaks, amplification in left side (red) and deletion in right side (blue). Each plot shows the focal level and arm level output, the green dashed line indicates the q-value threshold (q < 0.1). **b** The heatmap shows the mono or biallelic alteration of TP53 and CDKN2A.

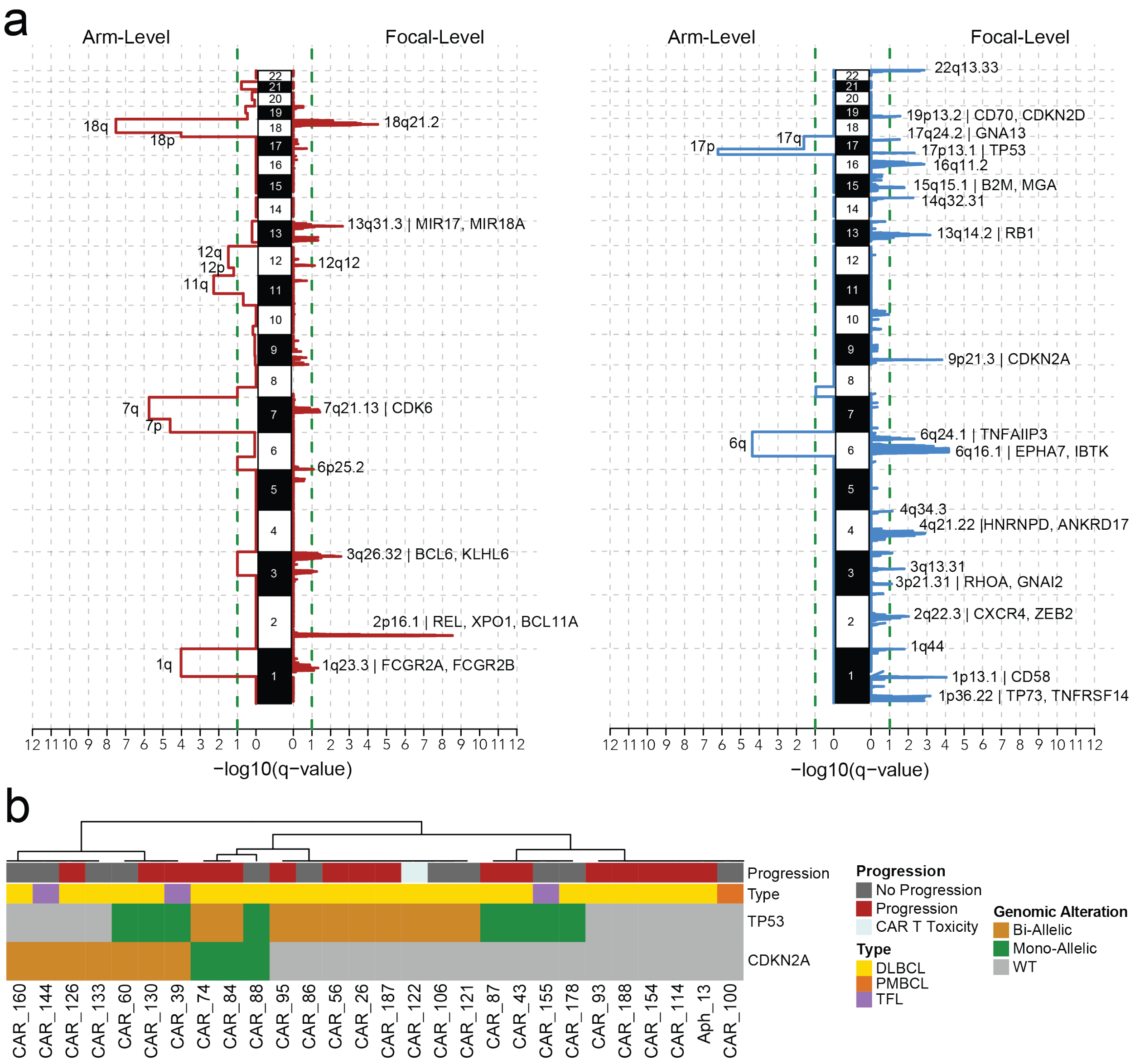

**Supplementary Figure 4. Overall survival Kaplan-Meier plots of statistically significant genomic alteration.**

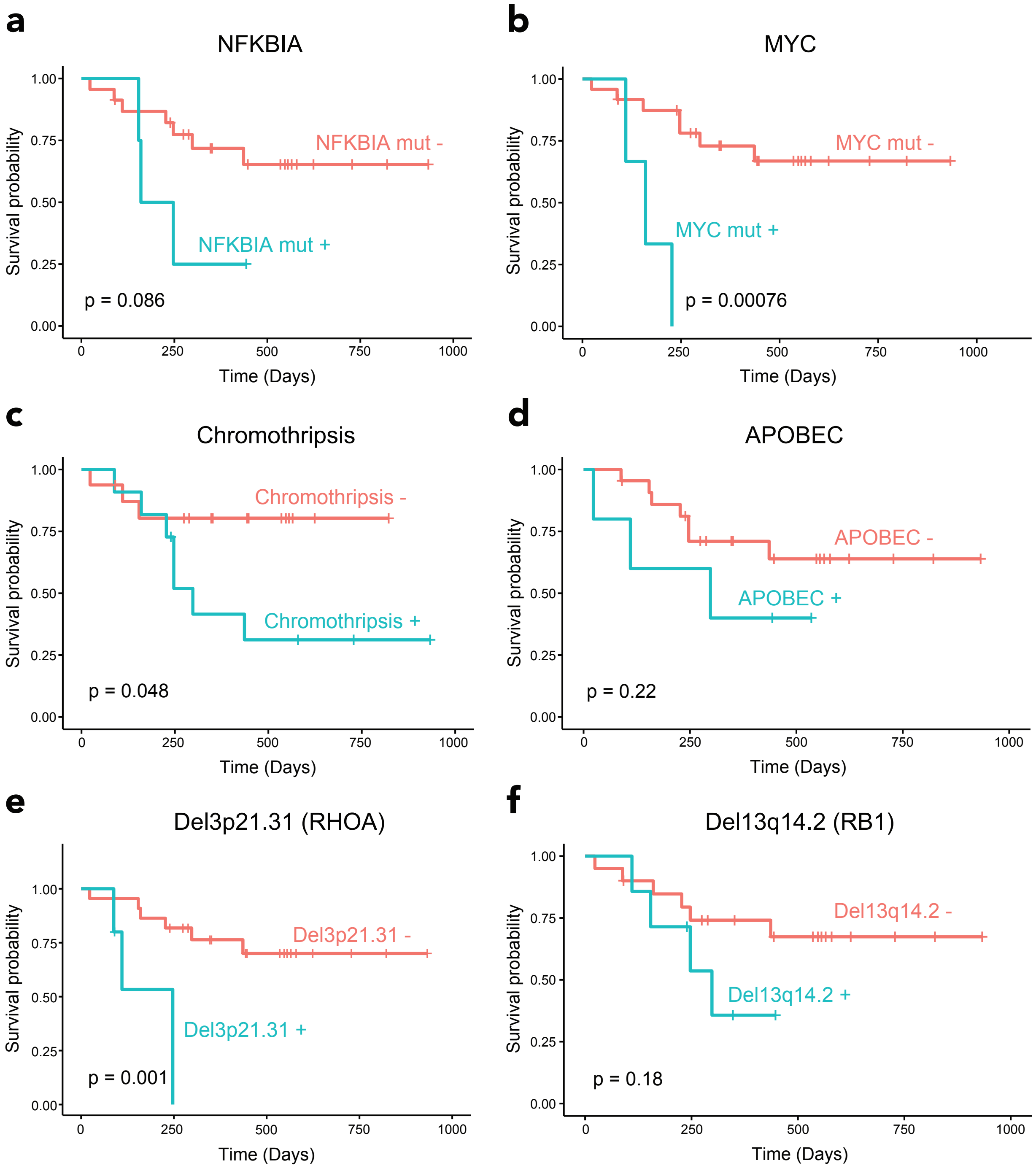

**Supplementary Figure 5. Kaplan-Meier curve of overall survival with the combination of statistically significant genomic alteration.**

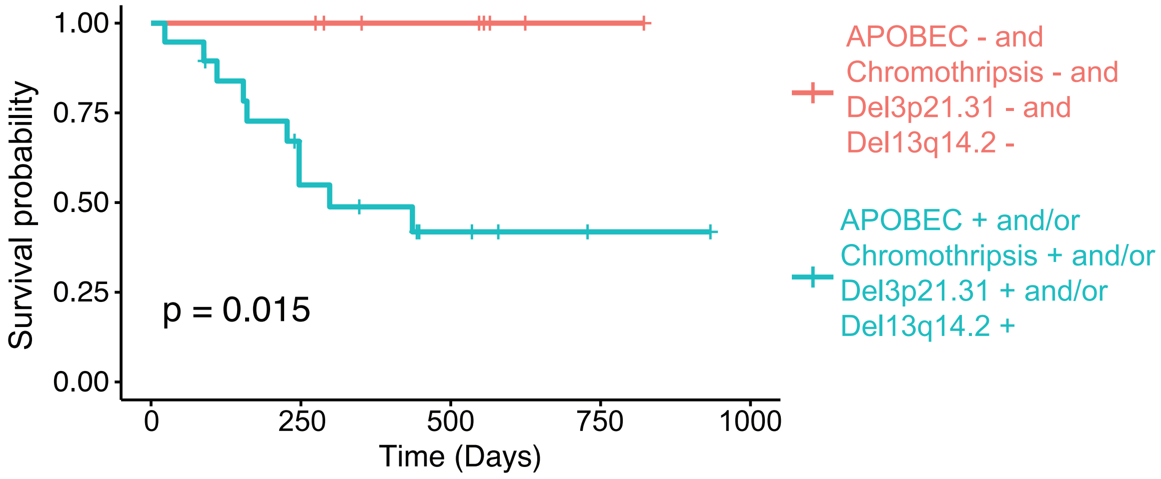

**Supplementary Figure 6. Statistically significant genomic alteration and association with INF signaling.** Enrichment plot from GSEA showing the enrichment of the T-cell INF signature (IFNG.GS) in patients without any significant genomic driver. The vertical black bars indicate the position of the genes in the signatures along the ranked gene list, the green line shows the enrichment score along the ranked gene list. The red to blue color bar shows the ranking of genes from up- to downregulated in patients with at least one genomic driver.

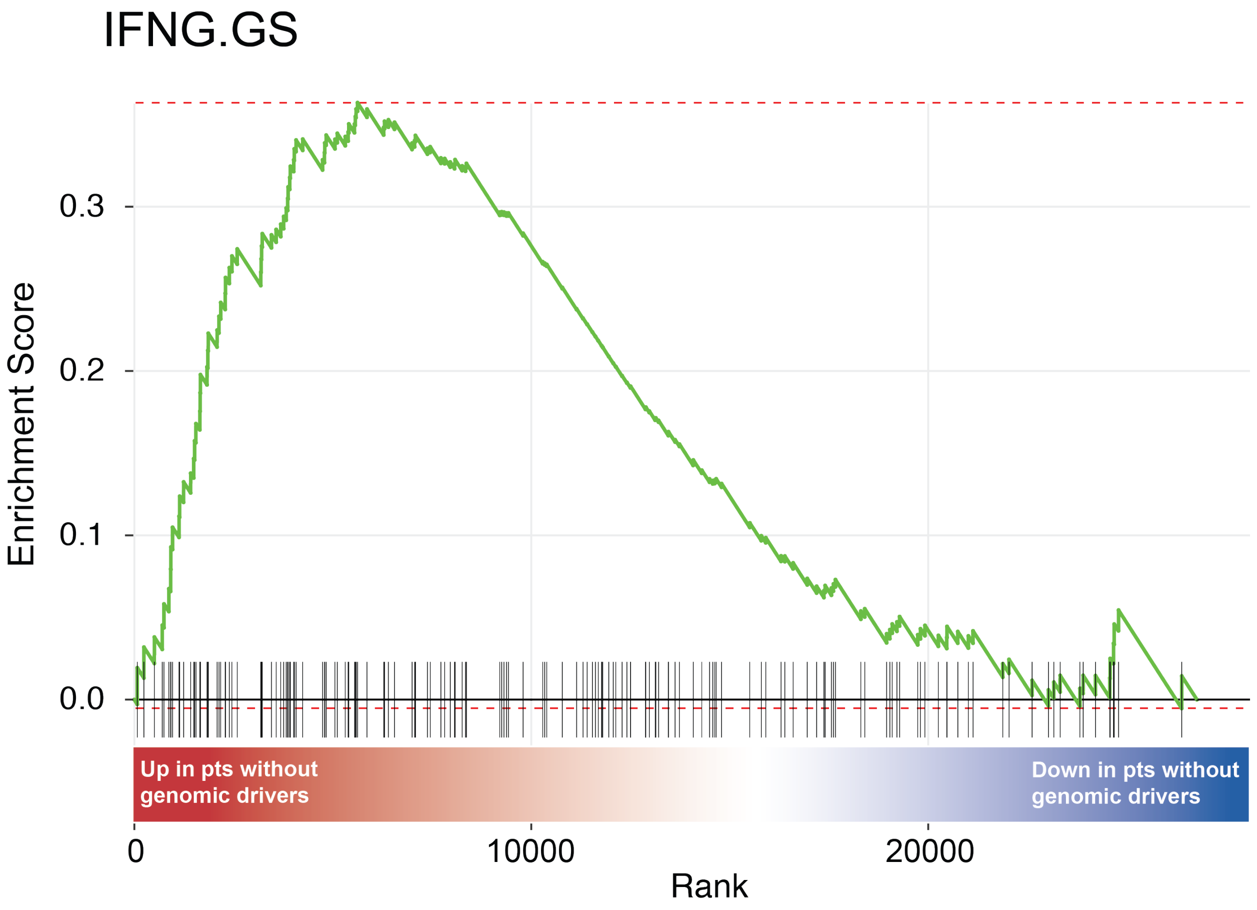

**Supplementary Table 1.** Coverage information.

| sample | median_coverage | type |
|---|---|---|
| **Aph_13** | 38.67 | normal |
| **Aph_13** | 42.58 | tumor |
| **CAR_26** | 51.11 | normal |
| **CAR_26** | 56.75 | tumor |
| **CAR_39** | 38.59 | normal |
| **CAR_39** | 50.08 | tumor |
| **CAR_39r** | 41.19 | tumor |
| **CAR_43** | 57.52 | normal |
| **CAR_43** | 46.26 | tumor |
| **CAR_56** | 36.87 | normal |
| **CAR_56** | 33.11 | tumor |
| **CAR_60** | 59.32 | normal |
| **CAR_60** | 37.39 | tumor |
| **CAR_65** | 48.86 | normal |
| **CAR_65** | 56.47 | tumor |
| **CAR_74** | 42.52 | normal |
| **CAR_74** | 43.6 | tumor |
| **CAR_78** | 45.31 | normal |
| **CAR_78** | 35.26 | tumor |
| **CAR_84** | 43.69 | normal |
| **CAR_84** | 50.65 | tumor |
| **CAR_84r** | 44.04 | tumor |
| **CAR_86** | 49.1 | normal |
| **CAR_86** | 46.77 | tumor |
| **CAR_87** | 35.63 | normal |
| **CAR_87** | 48.56 | tumor |
| **CAR_88** | 45.9 | normal |
| **CAR_88** | 39.61 | tumor |
| **CAR_91** | 40.79 | normal |
| **CAR_91** | 54.53 | tumor |
| **CAR_93** | 59.9 | normal |
| **CAR_93** | 39.26 | tumor |
| **CAR_95** | 66.23 | normal |
| **CAR_95** | 31.57 | tumor |
| **CAR_100** | 46.26 | normal |
| **CAR_100** | 47.52 | tumor |
| **CAR_106** | 41.83 | normal |
| **CAR_106** | 40.19 | tumor |
| **CAR_114** | 42.55 | normal |
| **CAR_114** | 42.76 | tumor |
| **CAR_121** | 34.17 | normal |
| **CAR_121** | 40.26 | tumor |
| **CAR_122** | 40.57 | normal |
| **CAR_122** | 30.39 | tumor |
| **CAR_126** | 42.4 | normal |
| **CAR_126** | 56.33 | tumor |
| **CAR_130** | 37.32 | normal |
| **CAR_130** | 34.39 | tumor |
| **CAR_133** | 45.21 | normal |
| **CAR_133** | 76.08 | tumor |
| **CAR_144** | 46.42 | normal |
| **CAR_144** | 44.25 | tumor |
| **CAR_154** | 48 | normal |
| **CAR_154** | 52.41 | tumor |
| **CAR_155** | 47.65 | normal |
| **CAR_155** | 35 | tumor |
| **CAR_160** | 41.16 | normal |
| **CAR_160** | 38.12 | tumor |
| **CAR_178** | 47.82 | normal |
| **CAR_178** | 46.43 | tumor |
| **CAR_187** | 45.72 | normal |
| **CAR_187** | 48.62 | tumor |
| **CAR_188** | 46.69 | normal |
| **CAR_188** | 44.38 | tumor |

**Supplementary Table 2.** Selected driver genes with dN/dScv algorithm (q < 0.1).

| gene_name | n_syn | n_mis | n_non | n_spl | n_ind | qglobal_cv |
|---|---|---|---|---|---|---|
| **TP53** | 0 | 24 | 0 | 2 | 1 | 0 |
| **KMT2D** | 0 | 6 | 14 | 2 | 10 | 0 |
| **TMSB4X** | 0 | 6 | 0 | 5 | 2 | 0 |
| **B2M** | 0 | 12 | 5 | 1 | 4 | 0 |
| **HLA-B** | 1 | 3 | 7 | 1 | 4 | 4.4611E-13 |
| **SOCS1** | 8 | 24 | 0 | 0 | 6 | 7.4352E-13 |
| **CD58** | 0 | 3 | 2 | 2 | 4 | 9.9451E-10 |
| **SGK1** | 4 | 19 | 0 | 5 | 1 | 1.1976E-07 |
| **CREBBP** | 0 | 11 | 5 | 2 | 3 | 3.4197E-07 |
| **TMEM30A** | 0 | 0 | 2 | 2 | 2 | 3.6802E-06 |
| **IGLL5** | 34 | 42 | 2 | 2 | 0 | 4.7593E-06 |
| **TNFRSF14** | 0 | 3 | 3 | 1 | 1 | 1.8452E-05 |
| **BCL2** | 23 | 34 | 0 | 0 | 0 | 0.00015909 |
| **GNAI2** | 0 | 6 | 0 | 0 | 1 | 0.00111564 |
| **MYD88** | 0 | 11 | 0 | 0 | 0 | 0.00127596 |
| **BTG2** | 3 | 14 | 0 | 0 | 1 | 0.00127596 |
| **CD79B** | 0 | 7 | 0 | 2 | 0 | 0.00287529 |
| **TNFAIP3** | 0 | 2 | 0 | 2 | 5 | 0.00290617 |
| **HLA-A** | 1 | 3 | 3 | 0 | 1 | 0.01010862 |
| **ITPKB** | 1 | 13 | 0 | 1 | 1 | 0.01207522 |
| **RPL22** | 0 | 0 | 0 | 0 | 3 | 0.01360407 |
| **OSBPL10** | 1 | 9 | 0 | 0 | 1 | 0.01942304 |
| **PIM1** | 14 | 19 | 1 | 1 | 2 | 0.02610203 |
| **FAS** | 0 | 4 | 0 | 0 | 2 | 0.02735161 |
| **ZFP36L1** | 2 | 8 | 1 | 0 | 4 | 0.0282139 |
| **ETV6** | 1 | 2 | 1 | 3 | 0 | 0.03612288 |
| **PRKCD** | 1 | 5 | 1 | 2 | 0 | 0.03612288 |
| **ARID1A** | 0 | 2 | 3 | 0 | 2 | 0.04390807 |
| **NFKBIE** | 0 | 2 | 0 | 1 | 4 | 0.04390807 |
| **GNA13** | 0 | 2 | 1 | 0 | 2 | 0.06081452 |
| **EBF1** | 1 | 4 | 0 | 4 | 0 | 0.06126379 |
| **HIST1H1C** | 2 | 16 | 0 | 0 | 1 | 0.06661659 |
| **BTK** | 0 | 4 | 0 | 1 | 1 | 0.09657792 |
| **NFKBIA** | 0 | 4 | 1 | 0 | 3 | 0.09657792 |
| **PTEN** | 0 | 1 | 1 | 1 | 1 | 0.09657792 |
| **RB1** | 0 | 0 | 0 | 2 | 2 | 0.09657792 |
